# Inflammation Drives Alternative First Exon usage to Regulate Immune Genes including a Novel Iron Regulated Isoform of Aim2

**DOI:** 10.1101/2020.07.06.190330

**Authors:** Elektra K. Robinson, Pratibha Jagannatha, Sergio Covarrubias, Matthew Cattle, Rojin Safavi, Ran Song, Kasthuribai Viswanathan, Barbara Shapleigh, Robin Abu-Shumays, Miten Jain, Suzanne M. Cloonan, Edward Wakeland, Mark Akeson, Angela N. Brooks, Susan Carpenter

**Affiliations:** Department of Molecular, Cell and Developmental Biology, University of California Santa Cruz, 1156 High St, Santa Cruz, CA 95064; Department of Biomolecular Engineering, University of California Santa Cruz; Department of Immunology, University of Texas Southwestern Medical Center, Dallas, TX, 75390-9093; Division of Pulmonary and Critical Care Medicine, Joan and Sanford I. Weill Department of Medicine, Weill Cornell Medicine, New York, NY 10021 USA

**Keywords:** Innate Immunity, Macrophages, Alternative Splicing, Aim2, 5’UTR, Nanopore Sequencing

## Abstract

Determining the layers of gene regulation within the innate immune response is critical to our understanding of the cellular responses to infection and dysregulation in disease. We identified a conserved mechanism of gene regulation in human and mouse via changes in alternative first exon (AFE) usage following inflammation, resulting in changes to isoform usage. Of these AFE events, we identified 50 unannotated transcription start sites (TSS) in mice using Oxford Nanopore native RNA sequencing, one of which is the cytosolic receptor for dsDNA and known inflammatory inducible gene, *Aim2*. We show that this unannotated AFE isoform of *Aim2* is the predominant isoform transcribed during inflammation and contains an iron-responsive element in its 5′UTR enabling mRNA translation to be regulated by iron levels. This work highlights the importance of examining alternative isoform changes and translational regulation in the innate immune response and uncovers novel regulatory mechanisms of *Aim2*.

**Summary Sentence:** Alternative first exon usage was the major splicing event observed in macrophages during inflammation, which resulted in the elucidation of a novel isoform and iron mediated regulatory mechanism of the protein coding gene, *Aim2*.

## Introduction

Macrophages are critical cells in the innate immune system that combat infection by initiating acute inflammatory responses. Acute inflammation is tightly coordinated and begins with the detection of pathogen-associated molecular patterns (PAMPs) by pattern recognition receptors (PRRs), which include Toll-Like Receptors (TLRs) (*1, 2*). These initial steps are followed by the activation of sequestered transcription factors, such as nuclear factor of kappa-B (NF-kB) and interferon regulatory factors (IRFs), which orchestrate pro-inflammatory and/or antiviral response signals involved in pathogen clearance (*2*). Once pathogens are cleared, macrophages express genes involved in the resolution of inflammation to return the host to homeostasis (*3*). Dysregulation of these pro-inflammatory pathways can have devastating consequences, leading to unresolved inflammation and chronic inflammatory diseases (*4*).

Recently the process of alternative splicing has emerged as another key mechanism by which the immune system is regulated. Alternative splicing is a regulated process enabling a single gene to produce many isoforms, thus increasing the complexity of gene function and the proteome (*2, 5*–*7*). Much of this occurs in a cell-type-specific and signal-induced manner (*8, 9*). Previous studies have shown that mouse and human macrophages exposed to inflammatory stimuli undergo alternative splicing (*2, 10*–*18*). Alternative splicing within the immune system can affect the type and magnitude of the inflammatory response, such as the production of a soluble form of TLR4 that is expressed upon LPS, which leads to inhibition of TNFα and NFκB serving as a negative feedback mechanism (*19, 20*). Additionally, this mechanism has been characterized within signaling molecules (*21, 22*), including TBK1 (*23*) and MyD88 (*24*), that produce the alternative RNA splice forms, TBK1s and MyD88s respectively, which function to limit the extent of the pro-inflammatory response. Alternative splicing can also result in the production of inflammatory signaling molecules, such as TRIF (*25*) and the proteins in the NFκB family(*9*) with altered activity or stability. Beyond changing the ORF of an mRNA molecule, elongating or shortening the first or last exon can impact post-transcriptional gene regulation and are important to consider when elucidating the regulatory mechanisms of immune genes (*26*–*28*), specifically underlying motifs in 5’UTRs (*29, 30*) and 3’UTRs (*31, 32*).

While inflammation-induced alternative splicing in both human and mouse macrophages has been investigated on a genome-wide scale (*2, 10*–*18*), to our knowledge, long read RNA-sequencing has not been utilized to investigate differential RNA isoform expression. Such an approach is necessary to fully appreciate the extent of alternative splicing as we know most transcriptome annotations are incomplete (*33*), and isoforms generated are cell-type and treatment specific (*33*–*36*).

Here we used both long and short-read RNA sequencing to uncover novel isoforms and classes of alternative splicing events following inflammation in human and murine macrophages. Interestingly the dominant conserved class of alternative splicing observed following inflammation is alternative first exon usage (AFE). AFE events can have a multi-level effect on protein diversity, regulating genes through alterations of the 5′UTR region, and directing the locality of proteins through alternative N-termini (*37*). We identified 50 unannotated AFE events in mice from native RNA sequencing, one of which is in the cytosolic receptor for dsDNA and known inflammatory inducible gene, *Aim2*. We show that this unannotated AFE isoform of *Aim2* is the predominant isoform produced during inflammation and contains an iron-responsive element in its 5′UTR, enabling mRNA translation to be controlled by iron levels. This work reveals that alternative splicing plays a crucial role in shaping the transient nature of the inflammatory response. Isoform expression is an additional layer of regulation within the immune response and therefore a possible contributing factor to the development of auto-immune and inflammatory diseases. Understanding the exact isoforms of genes that are expressed during an inflammatory response will enable us to design better targets for therapeutic intervention of these diseases.

## Results

### Global profiling of the cellular alternative splicing landscape in human and mouse macrophages post-inflammation

To identify alternative splicing events following inflammation, we performed whole transcriptome analysis on human monocyte derived macrophages (MDMs) and murine bone marrow derived macrophages (BMDMs) with and without lipopolysaccharide (LPS) treatment (Fig. 1A). We found that ∼50% of splicing changes (corrected p-value ≤ 0.25 and |ΔPSI| ≥ 10) were classified as alternative first exon (AFE) events following LPS activation in both human and murine macrophages (Fig. 1B, SFig. 1). We next identified 12 conserved AFE splicing events between human and mouse (SFig. 2). We validated the AFE changes upon stimulation on the already characterized *Ncoa7* (SFig. 3A-C) (*38, 39*) and *Rcan1* (SFig. 3D-F) (*40*), as well as a previously uncharacterized inflammatory specific isoform of *Ampd3* (SFig. 3G-I), in human and mouse primary macrophages using RT-PCR. Taken together, these results show the high prevalence and conservation of alternative first exons following inflammatory activation.

**Figure 1:**
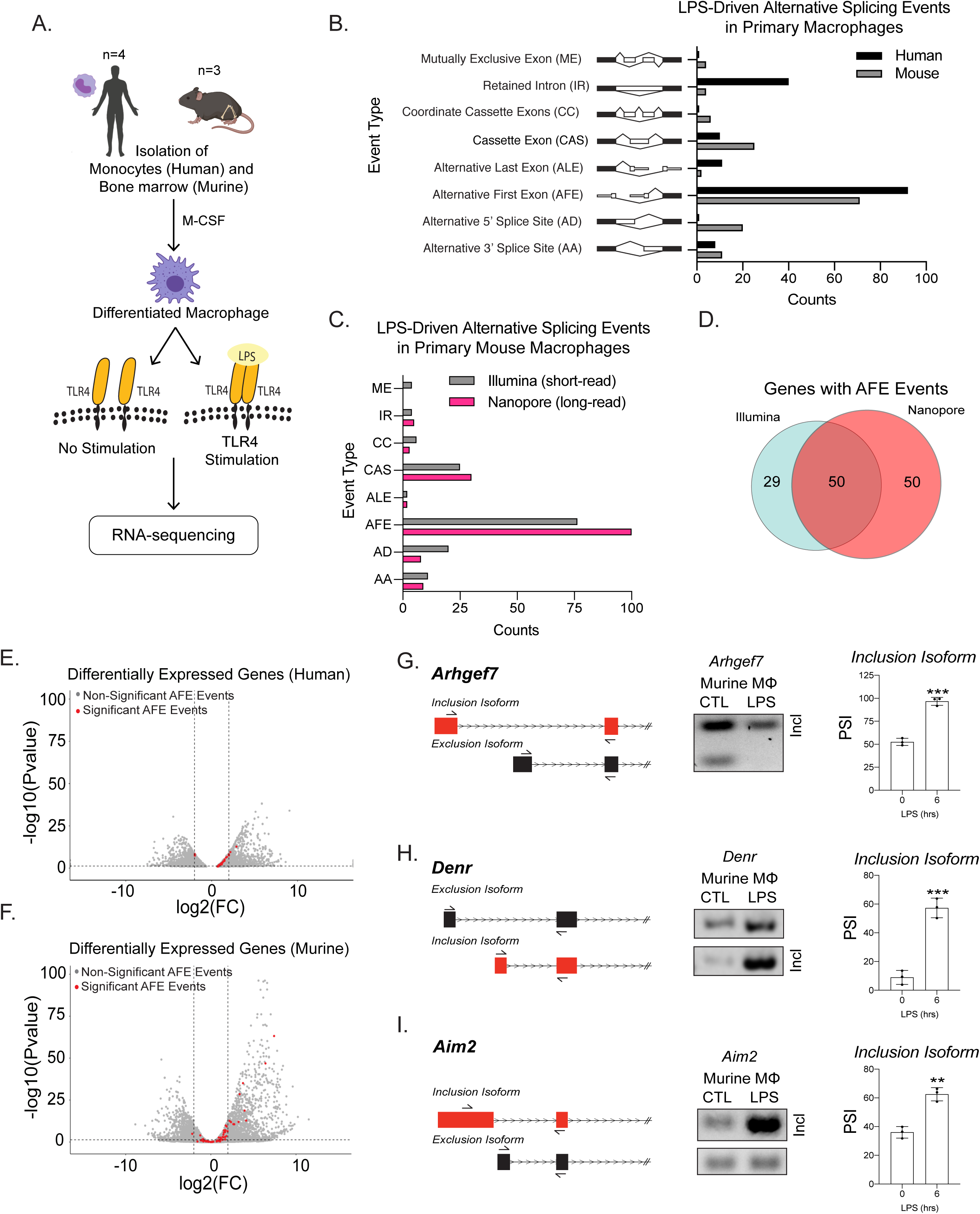
Global profiling of the cellular alternative splicing landscape in human and mouse macrophages post-inflammatory. **(A)** Diagram of RNA-seq library generation. **(B)** Categorization of significant splicing events in human and mouse macrophage. **(C)** Categorization of significant splicing events found in mouse BMDM +/− 6hrs LPS prior to and after incorporating high-confidence long-read isoforms identified using Nanopore sequencing. **(D)** Venn diagram representing unique and common genes with AFE events found in Nanopore or Illumina RNA-seq of primary BMDMs post-inflammatory stimulation. Volcano plots of all differentially expressed genes from RNA-seq of either human **(E)** or mouse **(F)** macrophages. Genes highlighted in red undergo significant AFE changes following inflammation. Schematic of AFE inclusion and exclusion isoforms, followed by RT-PCR gel results and PSI calculation for *Argehf7* **(G)** *Denr* **(H)** and *Aim2* **(I)**, were performed in biological triplicates, p-value assessed using student’s t-test.

**Figure 2:**
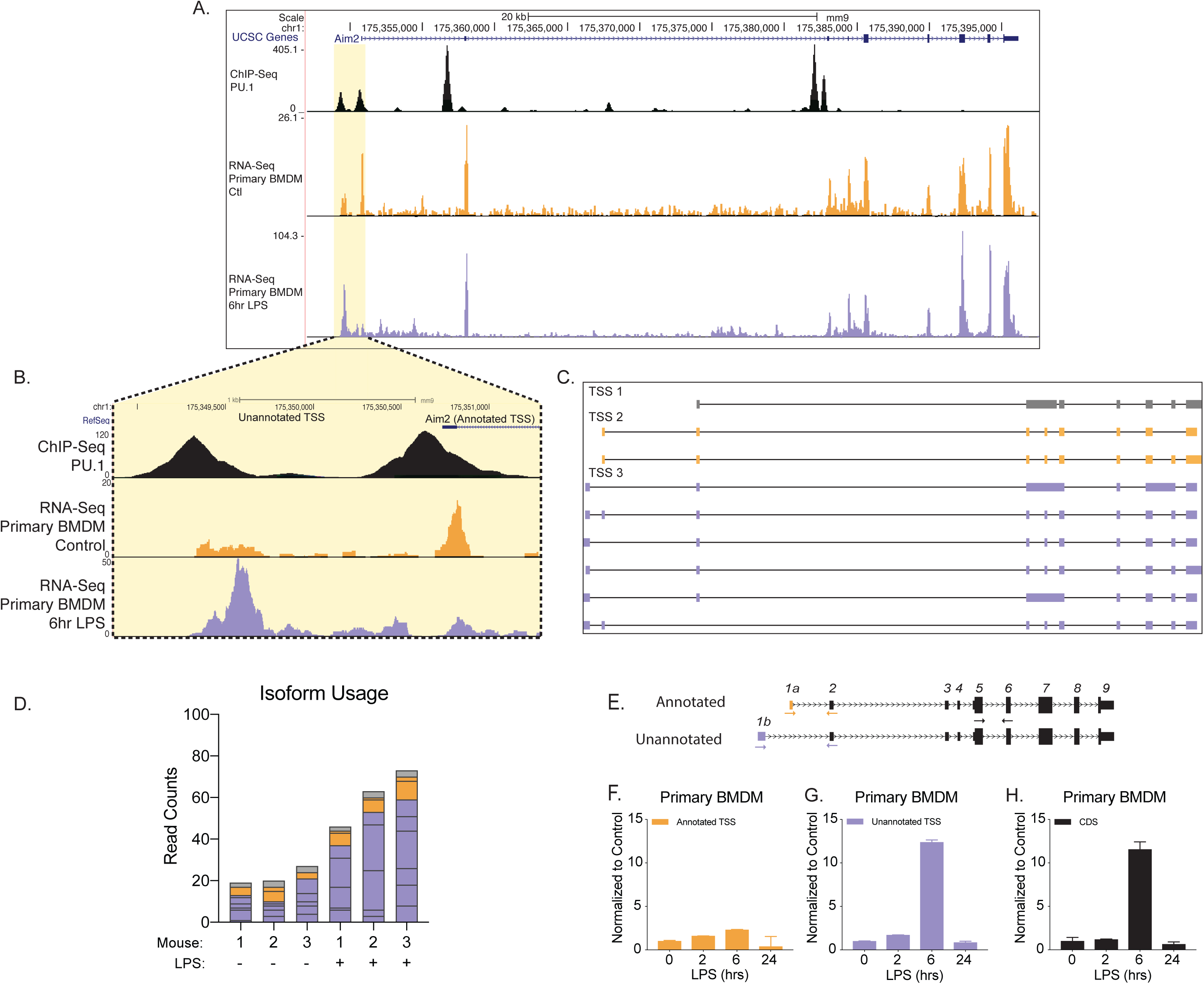
Identification of an unannotated promoter in *Aim2*. **(A-B)** The top track, in black, represents ChIP-seq data for a macrophage-specific transcription factor, PU.1. Peaks represent possible promoter regions; two distinct peaks of equal height are present at the annotated transcriptional start site for *Aim2* and about 1kb upstream of the transcriptional start site (TSS). The middle track, in orange, represents basal transcription in bone marrow-derived macrophages (BMDMs), while the bottom track, in purple, represents active transcription in BMDMs 6 hr LPS post treatment. **(C)** *Aim2* transcript isoforms identified in BMDMs by native RNA long-read sequencing through FLAIR analysis. Transcripts are categorized by promoter, denoted by grey, orange or purple. **(D)** The bar-chart represents data from long-read sequencing showing the abundance of each transcript isoform from BMDMs +/−6hrs LPS. **(E-H)** qRT-PCR was performed in biological triplicate, on primary BMDM RNA extracts that had been stimulated with LPS for indicated time points.

**Figure 3:**
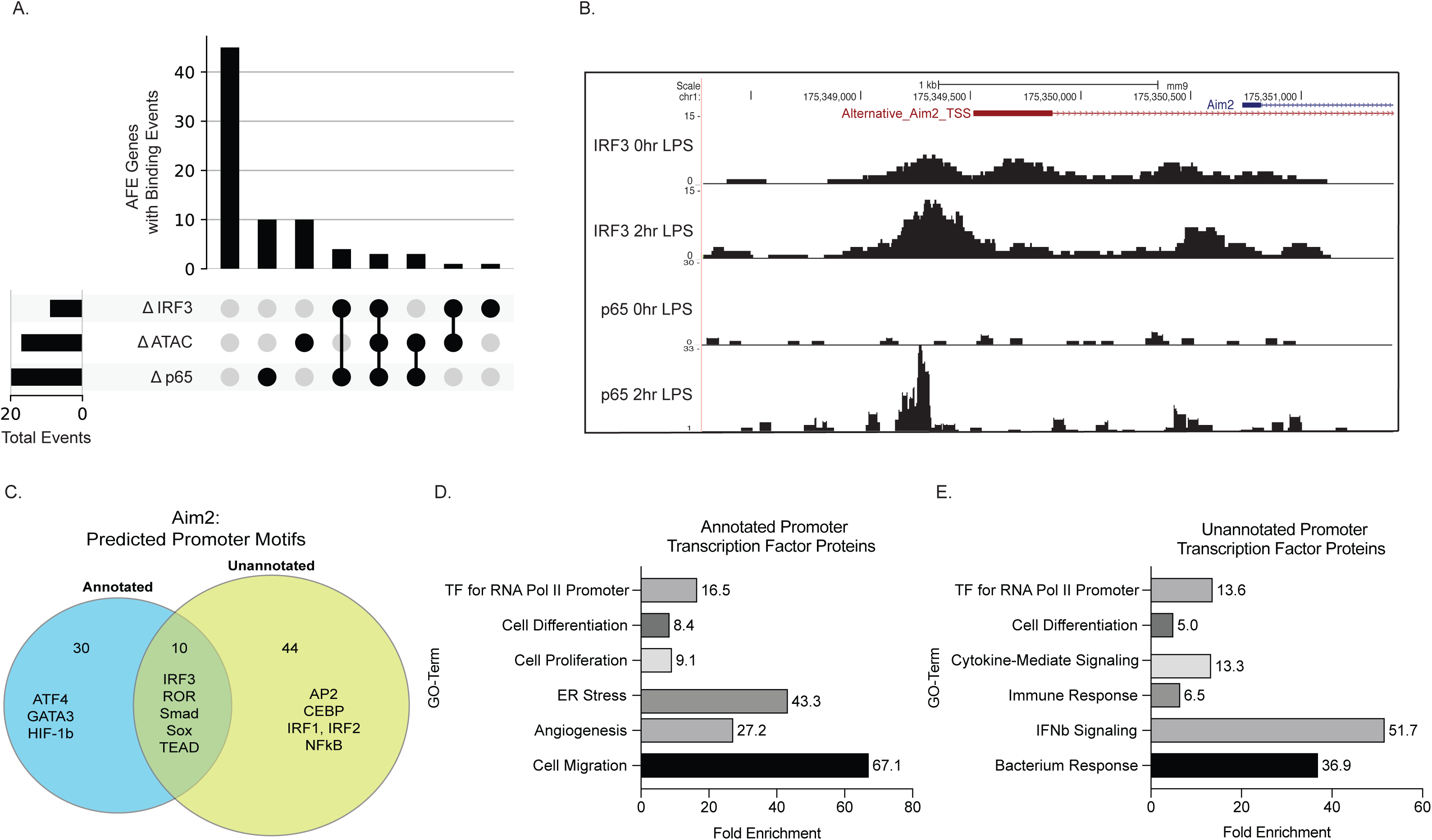
Novel inflammatory promoter of *Aim2* is regulated by IRF3 and p65. **(A)** UpSet plot showing the number of AFE events out of 77 total that have differential transcription factor binding and differential chromatin accessibility and all combinations of these sets. **(B)** mm9 genome browser shot between chr1:175,348,283-175,351,422 of ChIP-seq for IRF3 and p65 binding in BMDMs. **(C)** Venn diagram of all motifs defined using HOMER analysis within the annotated and unannotated promoter regions. **(D-E)** DAVID analysis examining the gene ontology of transcription factors at the annotated and unannotated promoters of *Aim2*.

A caveat to our analysis thus far was the reliance on annotated transcriptome assemblies. In order to determine if there are additional splicing events that are not captured using short-read sequencing (*33, 41*–*43*), we performed native RNA sequencing of murine macrophages with and without LPS treatment. We identified isoforms using Full-Length Alternative Isoform analysis of RNA (FLAIR) (*33, 44*) that also had promoter support identified from accessible chromatin (ATAC-seq) (*45, 46*). The FLAIR isoforms were then merged with the GENCODE M18 assembly (mm10) (*47*) to identify and quantify alternative splicing events (SFig. 4). Overall, the incorporation of long-read sequencing led to the identification of 50 novel and statistically significant AFE events that occur following inflammation (Fig. 1C-D). Interestingly, when identifying gene expression changes, we found that ∼50% of genes with AFE usage were not differentially expressed following inflammation (Fig. 1E-F) (Table 4-5), highlighting the importance of studying isoform usage for control of gene expression. Among the most statistically significant novel AFE first exon events were *Denr, Arhgef7*, and *Aim2*, which we validated using RT-qPCR (Fig. 1G-I) (SFig. 5).

**Figure 4:**
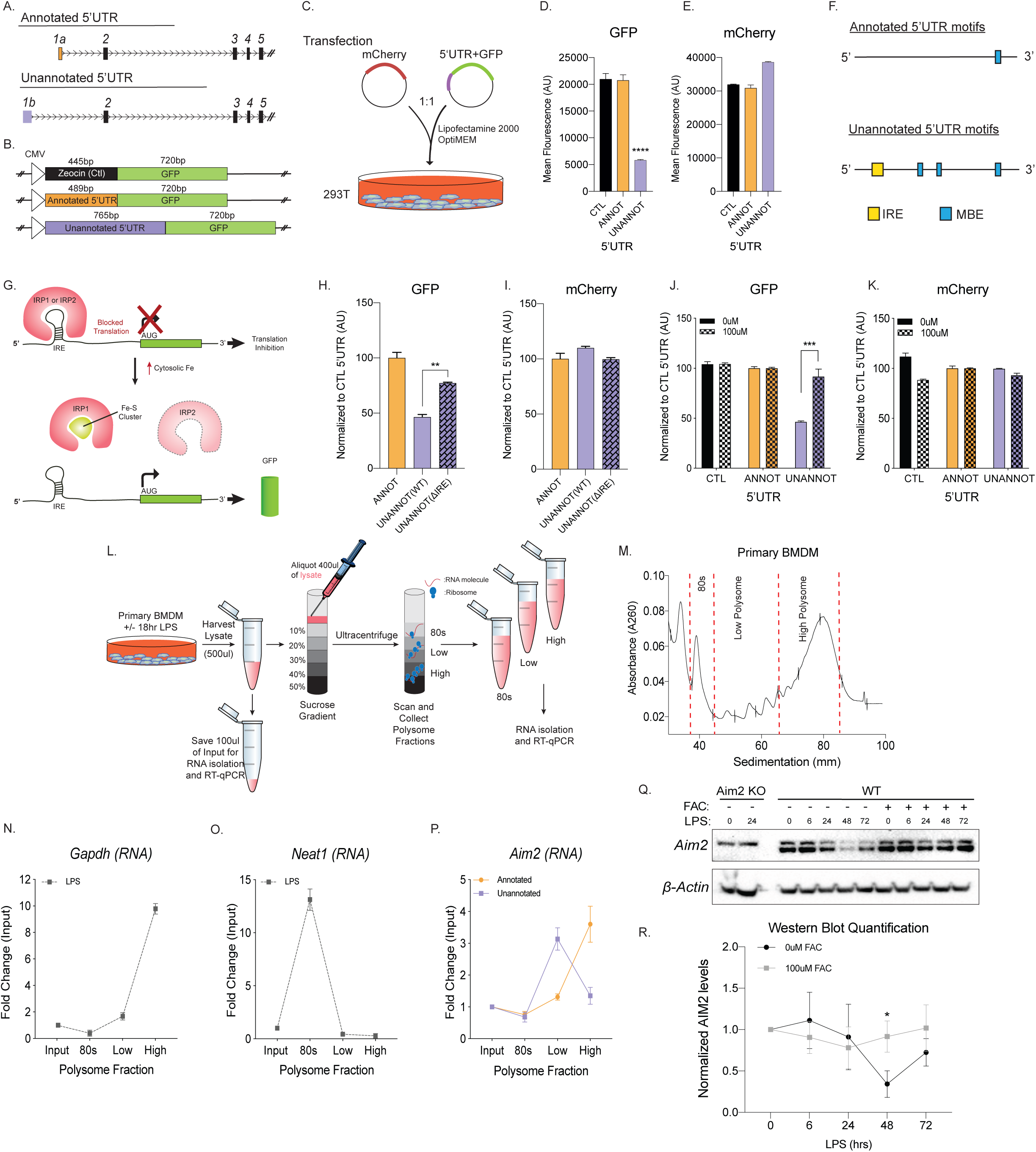
Unannotated 5′UTR of *Aim2* negatively regulates translation through a single iron responsive element. **(A)** Schematic of annotated and unannotated 5′UTR of most prevalent *Aim2* isoforms in mouse macrophages. **(B)** Diagram of cloning strategy of *Aim2’s* 5′UTR in GFP plasmid. **(C)** Transfection strategy of 5′UTR-GFP plasmids co-transfected with an mCherry control plasmid at a 1 to 1 ratio in 293T cells. **(D-E)** Flow cytometry of 293T cells 72 hr posttransfection with control annotated and unannotated 5′UTR of *Aim2* to measure GFP and mCherry (Ctl) protein fluorescence. **(F)** Using RegRNA2.0 a single iron-responsive element (IRE) was found in the alternative 5′UTR, in addition to multiple musashi binding elements (MBE). **(G)** Diagram of how an IRE functions in the cytoplasm of a cell within a 5′UTR. With low or normal levels of iron, Iron Binding Proteins (IRP1 or IRP2) bind to IRE elements and block translation. During high levels of iron, within a cell, IRP1 is sequestered by iron-sulfur (Fe-S) clusters and IRP2 is degraded, therefore allowing translation of the protein. **(H-I)** Flow cytometry of 293T cells 72 hr post-transfection of mCherry (Ctl), along with an annotated 5′UTR-GFP plasmid, unannotated 5′UTR-GFP plasmid, or a GFP plasmid containing the unannotated 5′UTR without the defined iron-responsive element (IRE). **(J-K)** Flow cytometry of 293T cells +/− 100uM ferric ammonium citrate (FAC) 72 hr post-transfection of mCherry (Ctl), along with an annotated 5′UTR-GFP plasmid or unannotated 5′UTR-GFP plasmid. **(L)** Overview of the polysome profiling protocol to analyze translation activity. **(M)** Cytoplasmic lysates from control and LPS treated cells were fractionated through sucrose gradients. Global RNA polysome profiles generated by the density gradient fractionation system are shown. A representative plot from stimulated primary BMDM fractionated samples is shown. The experiment was performed 4 times. **(N-P)** The relative distribution of *Gapdh* mRNA, encoding a housekeeping protein, *Neat1* long non-coding RNA (lncRNA) and *Aim2* mRNA were measured by RT-qPCR analysis of RNA using isoform-specific primer sets. Each of the gradient fractions are calculated as relative enrichment when compared to unfractionated input mRNA, standard deviation represents technical triplicate. **(Q)** Protein lysates of time course LPS stimulation of 0hr, 6hr, 24hr, 48hr and 72hr without and with 100uM of Ferric Ammonium Citrate (FAC, iron) added to immortalized WT BMDMs. Western blot performed on AIM2 and B-ACTIN. **(R)** Western blot quantification performed in FIJI, standard deviation represents biological triplicates, p-value assessed using student’s t-test.

### Identification of an unannotated promoter for Aim2

To better understand the potential functional consequence of AFE changes, we further examined the novel first exon event upregulated upon inflammatory activation in *Aim2. Aim2* is an interferon-stimulated gene (ISG), localized to the cytosol. *Aim2* is a dsDNA sensor, that upon recognition induces the formation of an inflammasome complex releasing IL1β and IL18 from the cell as a defense mechanism to control infection (*48*). Chromatin Immunoprecipitation (ChIP)-seq for the myeloid pioneering transcription factor PU.1 in primary BMDMs (Fig. 2A-B, top track, in black) (*49*) supported the presence of an additional promoter upstream of the RefSeq canonical isoform for *Aim2* (NM_001013779.2). Predominant isoforms (>=10% of total gene expression in a sample) assembled from native RNA sequencing with FLAIR identified the canonical RefSeq isoform and 5 unannotated isoforms that use the inflammatory activated promoter, revealing a new longer 5′UTR (Fig. 2C). Native RNA sequencing-based quantification provided additional support that the unannotated promoter usage is upregulated upon LPS stimulation. At steady-state, approximately 20% of reads map to *Aim2*’s transcript with the canonical promoter and 65% of reads map to transcripts with the upstream promoter, while following inflammatory activation, 14% of the reads map to transcripts with the canonical promoter and 81% of reads map to transcripts with the upstream promoter (Fig. 2D). To validate the change in *Aim2* AFE usage upon LPS stimulation, RT-qPCR was performed using exon spanning primers that were either specific to the annotated AFE or the unannotated AFE, in BMDMs (Fig. 2E). The expression profile of the annotated first exon is not induced by LPS stimulation (Fig. 2F), while the unannotated first exon and the CDS of *Aim2* are equally induced by LPS stimulation (Fig. 2G-H). Therefore, these data show that it is the novel isoform of *Aim2* is inflammatory-regulated and not the canonical isoform defined in RefSeq annotation, nor isoforms from the GENCODE annotation.

### The novel inflammatory promoter of Aim2 is regulated by IRF3 and p65

To gain insights into potential regulatory mechanisms controlling the expression of the 100 significant AFE events as well as *Aim2*, we assessed changes in chromatin accessibility during inflammatory activation in BMDMs. Analysis of ATAC-Seq (*46*) revealed differential peaks at the promoter regions for 25% of genes with significant AFE events suggesting that chromatin remodeling is one mechanism driving the expression of the AFE events (Fig. 3A). This mechanism is not what controls isoform usage for *Aim2*. The annotated promoter is accessible in all cells while the novel *Aim2* promoter is specific to myeloid progenitors and monocytes (*46, 50*) (SFig. 6A). In addition, the accessibility of both the annotated and unannotated promoters remain open despite the cell’s inflammatory status (SFig. 6B). Therefore, the expression of the new isoform is not due to chromatin remodeling of either promoter region (SFig. 6C).

Another mechanism that can drive AFE usage is transcription factor binding. We next analyzed ChIP-seq data of two major transcription factors that drive inflammation downstream of LPS, nuclear factor of kappa-B (NFkB, p65) and interferon response factor 3 (IRF3) (*46*). We found that p65 and IRF3 specifically account for another 25% of the AFE events, including the novel *Aim2* isoform, which we confirmed using multiple ChIP-seq data sets (Fig. 3A-B, SFig. 7AB). Further bioinformatic analysis of the two promoter regions driving the canonical and noncanonical isoforms of *Aim2* resulted in the identification of 106 individual transcription factor (TF) motifs within the annotated promoter and 121 motifs within the unannotated promoter (Table 8). Of these predicted motifs there were 54 motifs unique to the unannotated promoter (Fig. 3C). DAVID gene ontology (*51*) analysis of TFs specific to the annotated and unannotated promoter regions of *Aim2* confirms that the unannotated promoter is driven by inflammatory specific transcription factors including NF-κB and IRFs (Fig. 3D, E) (Table 9). Use of ATAC-seq and ChIP-seq for specific TFs has enabled us to determine the regulatory mechanisms driving 50% of the AFEs in our data. For the remaining 50%, there could be additional TFs, RNA binding proteins (*19*), or differential RNA stability driving their expression. Further work will be required to fully understand the complex regulation of all AFE events.

### Unannotated 5′UTR of Aim2 negatively regulates translation through a single iron responsive element

The novel inflammatory isoform identified here for *Aim2* acquired a longer 5′ untranslated region (UTR) compared to the canonical isoform (Fig. 4A). Previous studies have shown that longer 5′UTRs can affect the translation of a gene (*29, 52*). Using a GFP reporter system, the translational efficiency of the unannotated *Aim2* 5′UTR (767bp) was compared to the annotated 5′UTR (489bp) (Fig. 4B). The unannotated 5′UTR showed lowest mean GFP fluorescence units, suggestive of lower translational efficiency, as assessed by flow cytometry 72 hr post transient transfection in 293T cells (Fig. 4C-D), while equal mCherry fluorescence was observed for all co-transfected control constructs (Fig. 4E). To explore the mechanism of how the unannotated 5′UTR results in decreased translational efficiency, we used RegRNA2.0 to predict RNA regulatory motifs in the 5′UTRs (*53*). We identified a single iron-responsive element (IRE) within the unannotated 5′UTR, while Musashi binding elements (MBE) motifs were identified in both 5′UTRs (Fig. 4F) (Table 10). The finding that there are more structured motifs in the unannotated 5′UTR is also supported by the RNAfold Vienna package (*54*), which predicts the hairpin structure of the IRE element in the alternative 5′UTR (SFig. 8). Since the IRE motif is unique to the unannotated 5′UTR we hypothesized that this motif is critical in regulating translational efficiency.

When cells are at homeostasis, Iron Binding Proteins (IRP1/2) bind to IRE elements located within the 5’UTR (e.g. ferritin) can block translation, while IRE elements in the 3’UTR (e.g. transferrin receptor) can promote translation (*55, 56*). However, iron repletion results in the inactivation of IRP1/2 (*57*) (Fig. 4G). To experimentally test if the IRE motif within the unannotated 5′UTR acts as a translational repressor, we removed the element using site-directed mutagenesis which led to an increase in GFP expression by ∼20% compared to the annotated 5′UTR (Fig. 4H-I). Next we exogenously added 100 uM ferric ammonium citrate (FAC) to overload the cells with iron and determined if this can rescue the observed decrease in translational efficiency from our unannotated 5′UTR. Upon FAC administration, the relative GFP expression of the unannotated 5′UTR plasmid increased by ∼50%, while mCherry control was unchanged, suggesting that the translational efficiency of the unannotated 5′UTR can be rescued with iron supplementation (Fig. 4J-K). From these results, we conclude that the predicted IRE motif within the unannotated 5′UTR of *Aim2* functions as an IRE to control translation.

To test if the IRE motif in the unannotated 5′UTR of *Aim2* acts as a translational regulator endogenously, we performed polysome profiling followed by RT-qPCR on primary BMDMs in the presence and absence of LPS for 18 hr to determine the translational competency of the isoforms of *Aim2* (Fig. 4L-M). The relative distribution of our positive control gene, *Gapdh*, which encodes a highly expressed housekeeping protein, is enriched in the high polysome fraction as expected (Fig. 4N). As a negative control, we examined Neat1, a long non-coding RNA (lncRNA) that is not detected in polysomes, nor translated (Fig. 4O). Using isoform-specific primer sets (Fig. 2E) for the annotated and unannotated *Aim2* isoforms, we find that the annotated isoform is enriched in the high polysome fraction during no treatment, while the novel isoform is enriched in the low polysome fraction upon inflammatory stimulation (Fig. 4P). These data show that the unannotated *Aim2* isoform has a lower translational efficiency compared to the canonical form. Finally, to further validate the effect of expression of the unannotated *Aim2* isoform on the level of protein, we performed a time-course LPS stimulation for 72 hr, with and without the treatment of iron (FAC) and measured AIM2 expression by western blot. AIM2 is expressed basally and significantly decreases upon LPS treatment at the 48 hr time point, most likely as a control mechanism to return the pathway to homeostasis and limit the inflammatory stimulation. When FAC is added to cells, AIM2 expression does not decrease at the 48 hr time point suggesting that it is indeed the IRE element that is driving this decrease in AIM2 observed in the wild type cells (Fig. 4Q-R).

## Discussion

While we have come a long way in determining the transcriptomes of immune genes to better understand signaling pathways, very little work has focused on the role that mRNA isoforms play (*2, 58, 59*). Over 95% of genes have more than one mRNA transcript due to alternative splicing but the regulatory importance of these splicing events are not fully understood (*60*–*62*). On a gene-by-gene approach, alternative splicing has been shown to play a role in health and disease by shaping the proteome (*63*–*65*). Globally, a number of labs have tackled the prevalence of alternative splicing *in vitro* and *in vivo* showing that alternative splicing can affect both the nature and duration of inflammation (*10, 11, 14*). To date no one has examined conservation of these mechanisms using primary cells or utilized long read sequencing to build the transcriptome *de novo* to obtain a complete understanding of the extent of alternatively expressed isoforms generated following an immune response.

In our study, we demonstrate a conservation of splicing, specifically alternative first exon (AFE) events, in both human and murine macrophages. We found that there are 12 genes that have AFE in both human and mouse macrophages (SFig. 2). Most studies to date have focused on isoform changes linked to genes that are differentially expressed following inflammation and interestingly these 12 genes would have been previously overlooked because many of them are not differentially expressed emphasizing the importance in studying isoform expression in all conditions (Fig. 1E-F). Of the 12 conserved genes, 7 AFE isoforms have been previously studied in some context including: Rps6ka1 (*66*), Ncoa7 (*38, 39*), Rcan1 (*40*), Wars (*67, 68*), Arap1 (*69*), Adar (*70, 71*) and Sgk1 (*72*–*74*). While this validates our technique, it is important to note that none of these genes had been connected to inflammation and formally shown to be conserved mechanisms of regulation, besides Wars (*67, 68*). This also highlights our method’s ability to accurately identify inflammatory regulated RNA isoforms, in addition to the uncharacterized AFE events of Cept1, Ampd3, Snx10, Elf1 and Tspan4. Furthermore, Snx10 and Elf1, two proteins studied outside the context of inflammation, have been implicated in chronic inflammatory disease and our study may suggest new insights into how alternative splicing could be regulating these genes (*64, 65*). We further validated Ncoa7, Rcan1 and Ampd3 in human and murine macrophages using RT-PCR (SFig. 3).

To overcome the current limitations of any transcriptome build we used direct RNA nanopore technology on primary murine macrophage nanopore to build our own transcriptome *de novo* with the aim of identifying novel full transcriptional isoforms (*33, 44*). Following this, we identified hundreds of novel isoforms resulting in 50 novel AFE events (Fig. 1), including an unannotated mRNA isoform of the well-studied gene protein absent in melanoma 2 (*Aim2*).

*Aim2* is characterized as an interferon inducible gene (*75*) (PMID: 10454530), functioning as a cytoplasmic dsDNA sensor leading to the formation of an inflammasome and eventual cleavage and release of pro-inflammatory cytokines of IL1β and IL18 (*76*–*78*). Our study highlights that it is an alternative mRNA isoform of Aim2 that is inducible, and that this upregulated transcript is translated less efficiently compared to the canonical isoform. This novel finding goes against the existing assumption that induced gene expression results in induced protein expression. (Fig. 1I, Fig. 2, Fig. 4). FLAIR-identified transcripts (*33*) show three clear transcriptional start sites (TSS) for *Aim2* and only one of those TSSs is inflammatory inducible (Fig. 2C-D). RT-qPCR further solidifies that the annotated TSS of *Aim2* is not inducible, while the unannotated TSS of *Aim2* is LPS inducible (Fig. 2E-H). This result highlights the need for cell-type specific transcriptome annotations if one is to have a complete understanding of the transcriptome and proteome of a given cell.

We further investigated what drove the expression of this new *Aim2* isoform, as well as what drove the expression of all the AFE genes. Using ATAC-seq (*46, 50*) and ChIP-seq (*46*) data sets we were able to determine that the AFE events were driven partially by chromatin accessibility and inflammatory specific transcription factors, while 50% were unaccounted for (Fig. 3A). Further analysis incorporating more transcription factors will be necessary to determine the regulatory mechanism. Interestingly, the annotated *Aim2* promoter accessibility is constitutively open across all hematopoietic cells, while the unannotated *Aim2* promoter is only accessible in myeloid progenitors or terminally differentiated cells (SFig. 6A), meaning that the novel mechanism of *Aim2* regulation is specific to myeloid cells only. Furthermore, *Aim2* annotated and unannotated promoter usage is not driven by chromatin accessibility (SFig. 6B-C) but is solely driven by the activation of inflammatory specific transcription factors (Fig. 3B-E).

There is no difference in the open reading frame of the novel isoform of Aim2 when compared to the annotated transcript, therefore we turned our attention to a possible regulatory mechanism within the 5′UTR (*79*). Broadly, UTRs play crucial roles in the post-transcriptional regulation of gene expression, including alteration of the mRNA translational efficiency (*80*), subcellular localization (*81*) and stability (*82*). Post-transcriptional regulatory mechanisms of Aim2 have not been previously studied. Using RegRNA2.0 (*53*) we identified an Iron Responsive Element (IRE), a unique regulatory motif within the novel 5′UTR of Aim2 (Fig. 4F). Utilizing a GFP reporter plasmid, we were able to determine that the IRE motif was functional, by recapitulating the same experiments used on the protein Ferritin, the first functional IRE motif ever studied (*83*). Finally, we showed the inflammatory specific mechanism regulating AIM2 protein expression by performing a western blot on primary BMDMs with and without Ferric Ammonium Citrate (FAC) during a 72 hr LPS time-course experiment. AIM2 protein is basally expressed and while the transcript is inducible, specifically the novel isoform we identify here we do not observe an increase in expression of AIM2 protein by western blot. In fact, we find that AIM2 protein decreases following inflammation and this can be reversed by iron supplementation. This could be a critical regulatory step that has evolved to ensure the AIM2 pathway is switched off following its formation and activation of the inflammasome.

These results demonstrate that the inflammatory specific mRNA isoform of *Aim2* has lower translational efficiency than the canonical form and that protein translation can be increased by the addition of iron. Crane *et al*. (*84*) demonstrate that ROS can contribute to activation of AIM2 inflammasome in mouse macrophages. Our proposed mechanism of translational regulation of AIM2, is through an Iron Responsive Element (IRE), which is known to directly interact with IRP proteins (*55*–*57*). Interestingly, IRP2 degradation can be driven not only through iron, but also through ROS and RNS (*85*), further supporting this novel IRE regulatory mechanism of AIM2 protein expression. Finally, Cheng *et al*. has shown that AIM2 is regulated by oxidative stress and show that over activation of AIM2 inflammasome can contribute to pancreatic tumorigenesis, all within the environment of mitochondrial iron overload (*86*). This newly identified isoform, with an IRE specific translational mechanism provides mechanistic understanding to these recent studies of Aim2 (*84, 86*). These findings could have significance for better understanding the mechanisms driving pathology in inflammatory disease such as systemic lupus erythematosus (SLE) (*87*). AIM2 expression levels have been correlated with severity of inflammation in SLE patients (*88*) and it is well known that iron is dysregulated in this disease (*89*). It is possible that AIM2 levels remain high in SLE patients due to dysregulated iron; therefore, homeostasis in macrophages cannot be maintained.

In summary, signaling within macrophages has the ability to fight infection but also contribute to pathological inflammation associated with a wide variety of diseases. While there are multiple regulatory checkpoints in place to control inflammation, we propose that alternative splicing and translational regulation play critical roles in maintaining this type of control. A better understanding of the molecular mechanisms that control inflammatory-regulated genes, including *Aim2*, could provide new targets for therapeutic intervention of autoimmune and inflammatory diseases.

## Supporting information

Table1_Human_Illumina

Table2_Mouse_Illumina

Table3_Mse_Illumina_Nanopore

Table4_DESeq2_Hu

Table5_DESeq2_Mse

Table6_Differential_ATACseq

Table7_p65_irf3_peak

Table8_Mse_Homer_Aim2

Table9_Promoter_DAVID

Table10_RegRNA_Aim2

## Acknowledgements

We thank Eric Martin for sharing his time and knowledge on HOMER. We would also like to thank Alison Tang with sharing her knowledge and assisting with FLAIR isoform analysis. Finally, we would like to thank Kevin S. Johnson and Karen Ottemann who lent reagents for the IRE mutagenesis experiments.

## Author contributions

EKR performed all the molecular biology experiments from Fig 1, 2 and 4. PJ carried out all the bioinformatic analyses on alternative splicing from both mouse and human LPS stimulated macrophages. SCo assisted in designing of cloning experiments and generating samples for polysome profiling in Fig 4. MC performed promoter and AFE analysis in Fig 3 and EKR performed HOMER promoter analysis on Aim2 promoters. BS cultured the primary mouse macrophages and generated the illumina, while RAS generated the Nanopore RNA-sequencing libraries. RS analyzed the Nanopore sequencing datasets, with assistance from MJ. RS performed the human primary macrophage culturing and sequencing experiences. SMC assisted in the construction of the Iron related experiments and background knowledge in the field. ANB, SCa, EKR and PJ conceived and coordinated the project. EKR, PJ, ANB and SCa wrote the manuscript with input from all other co-authors.

## Data and Materials Availability

All data are available in the manuscript or the supplementary materials. RNA-seq raw data have been deposited in the NCBI Gene Expression Omnibus database under accession numbers GSE141754.

## Materials and Methods

### Human PBMC derived macrophage differentiation and in vitro stimulation

Human peripheral blood mononuclear cells (PBMC) were enriched by density gradient centrifugation of peripheral blood from healthy human donors through a Ficoll-Paque PLUS (GE Healthcare) gradient. Monocytes were isolated from PBMC by negative selection using the EasySep™ Human Monocyte Isolation Kit (STEMCELL Technologies) according to the manufacturer’s instructions. To differentiate monocytes into macrophages, recombinant human M-CSF (50ng/mL) was used in RPMI-1640 medium with 10% FBS, 2mM L-glutamine, 10mM HEPES, 1mM sodium pyruvate, 100U/ml penicillin, and 100 µg/mL streptomycin. The culture medium which contained fresh recombinant human M-CSF was replaced every 2 days.

### Cell culture, Mouse Macrophage Differentiation and Stimulation

Cells were cultured in D-MEM with 10% fetal bovine serum (FCS) supplemented with penicillin/streptomycin or ciprofloxacin. Primary BMDM were generated by cultivating erythrocyte-depleted bone marrow cells in the presence of 30% L929 supernatant and the cells were used for experiments 6-9 days after differentiation. J2Cre virus (Blasi et al., 1989) was used on day 3/4 after isolation of bone marrow cells to establish transformed BMDM cell lines. BMDMs were cultivated in the presence of J2Cre virus for 48h and L929 was then gradually tapered off over 6-10 weeks depending on the growth pattern of transformed cells.

### In vitro stimulation of macrophages

Bone derived macrophage cells were primed with 100uM of Ammonium Ferric Citrate (FAC) for 24hrs prior to TLR stimulation. Bone marrow derived macrophage cells were stimulated with Toll-like receptor (TLR) ligands for the indicated time points using the following concentrations: Lipopolysaccharide (LPS) 100ng/ml (TLR4). For RNA and protein isolation, 1-2×10^6^ cells were seeded in 12-well format.

### Cloning Strategy for 5’UTR GFP Plasmid

The GFP reporter plasmid was CMV-Zeo-t2A-GFP. Zeocin is flanked by NheI and AgeI. The sequence of the annotated and unannotated 5’UTR were used as defined by the UCSC RefSeq and our sequencing results to be. Using KAPA HiFi HotStart ReadyMix PCR Kit (Kapa Biosystems) the two 5’UTRs were amplified from cDNA.

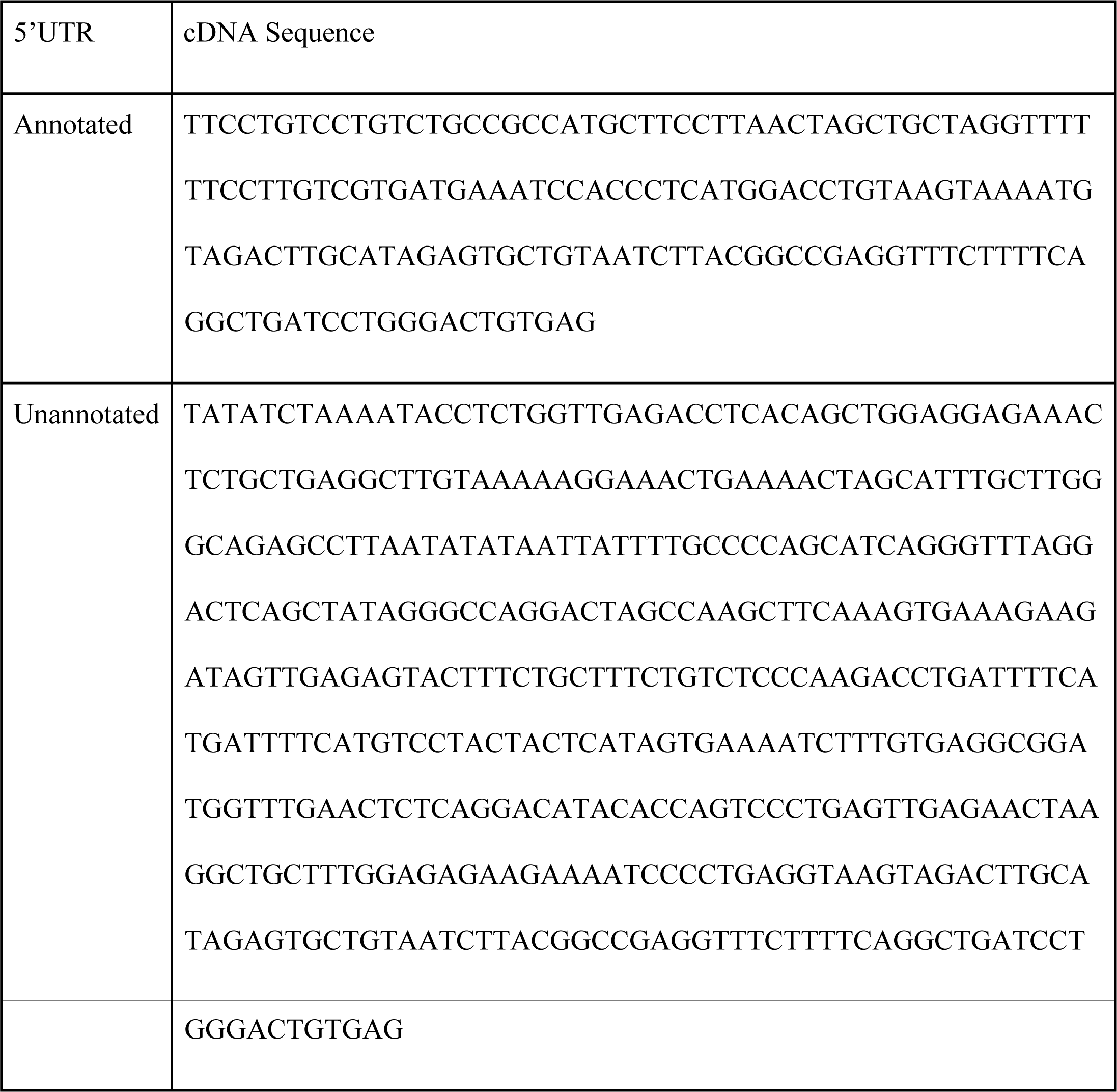

Primers:

*cDNA_F_Annotated:* TTCCTGTCCTGTCTGCCG

*cDNA_F_Unannotated:* TATATCTAAAATACCTCTGGTTGAGACCTC

*cDNA_R_5’UTR:* CTCACAGTCCCAGGATCAGC

The 5’UTRs were then PCR amplified with primers containing restriction enzyme sites for AgeI and NheI.

*NheI_F_Annotated:* GGT**GCTAGC**TTCCTGTCCTGTCTGCC

*NheI_F_Unannotated:* GGTGGT**GCTAGC**TATATCTAAAATACCTCTGGTTGAGA

*AgeI_R_5’UTR:* GGT**ACCGGT**CTCACAGTCCCAGGATCAGC

PCR products, as well as the GFP plasmid, were then processed using AgeI and NheI restriction enzymes overnight. These samples were run on a 1% agarose gel. A gel extraction was completed for each band using the PCR clean-up Gel extraction kit (Machery-nagel). The PCR product was confirmed using Sanger Sequencing.

### Site Directed Mutagenesis

Set up PCR reaction with 1ul 279 Plasmid, with unannotated Aim2 5’UTR, 5ul 10x Phu buffer, 1ul F primer (0.1ug/ul) [Remove_IRE_F − CCCTGATGCTGGGGCAAAATAATTATAAATGCTAGTTTTCAGTTTC], 1ul R primer (0.1ug/ul) [Remove_IRE_R − GAAACTGAAAACTAGCATTTATAATTATTTTGCCCCAGCATCAGGG], 1ul dNTP (10nM), 1ul Phu polymerase and 40ul dH2O. PCR program: 95°C 1min, 18 cycles of 95°C 30sec, 55°C 1min, 72°C 1min, then end PCR with 72°C 1min and 4°C hold. Add 0.5ul of Dpn1 (NEB) to 25ul PCR reaction. Incubate at 37 degrees celsius for 1hr to digest parental DNA. Transform digested and undigested plasmid into DH5α competent cells. Pick ∼10-15 colonies and start overnight cultures. Colony PCR plasmids using Dpn1_Colony_PCR_F − TTGGCTAGTCCTGGCCCTAT and Dpn1_Colony_PCR_R − GCTGGTTTAGTGAACCGTCAG to check for 20bp deletion on a 3% Agarose Gel. Grow up colonies that have deletion, miniprep plasmids and send to sequetech for sequence verification.

### Transfection of 5’UTR GFP and mCherry Plasmid

A 1:1 ratio of the GFP vector containing the mature sequence of Aim2 5’UTR (annotated or unannotated) or zeocin and a plasmid containing mCherry were transfected into 293T cells for 48-72hrs. A 6-well plate of HEK293Ts were plated the night before with a concentration of 2×10^5^. HEK293Ts cells were primed with 100uM of Ammonium Ferric Citrate (FAC) for 24hrs prior to transfection. Transfection was performed on HEK293Ts (+/− 100uM FAC) using lipofectamine 2000, serum free OPTI-MEM media was used as a transfection reagent according to manufacturer’s instructions, and a (1:1) concentration of the 5’UTR GFP reporter plasmid and the mCherry control plasmid. HEK293Ts were visualized via flow cytometry 48-72hrs post transfection.

### Maintenance of mice

UCSC and the Institutional Animal Care and Use Committee maintained mice under specific pathogen-free conditions in the animal facilities of ‘University of California Santa Cruz (UCSC) in accordance with the guidelines.

### Polyribosome Profiling

Prior to lysis, cells were treated with cycloheximide (100 mg/mL), 10 min at 37°C 5%CO2. Cells were washed three times with ice cold PBS and lysed in ice cold buffer A (0.5% NP40, 20 mM Tris HCl pH 7.5, 100 mM KCl and 10 mM MgCl2). Lysates were passed three times through a 23G needle and incubated on ice for 7 min. Extracts were then centrifuged at 10K rpm for 7 min at 4°C. The supernatant was collected as crude cytosolic extract. Cytosolic extracts were overlaid on 10%–50% sucrose gradients prepared in 20 mM Tris HCl pH 7.5, 100 mM KCl and 10 mM MgCl2 buffer (prepared using the Gradient Station, Biocomp Instruments). Gradients were then ultracentrifuged at 40K rpm for 1h 20 min at 4°C using an SW41 in a Beckman ultracentrifuge. Individual polyribosome fractions were subsequently purified using a Gradient Station (Biocomp Instruments) and stored in (1:3) TRI Reagent.

### RNA isolation, RT-qPCR

Total cellular RNA from BMDM cell lines or tissues was isolated using the Direct-zol™ RNA MiniPrep Kit (Zymo Research) according to manufacturer’s instructions. RNA was quantified and controlled for purity with a nanodrop spectrometer. (Thermo Fisher). For RT-qPCR, 500-1000 ng were reversely transcribed (iScript Reverse Transcription Supermix, Biorad) followed by RT-PCR (iQ SYBRgreen Supermix, Biorad) using the cycling conditions as follows: 50°C for 2 min, 95°C for 2 min followed by 40 cycles of 95°C for 15 sec, 60°C for 30 sec and 72°C for 45 sec. The melting curve was graphically analyzed to control for nonspecific amplification reactions. Quantitative RT-PCR analysis was performed with the following primers listed below:

*Mse_Aim2_F_Annotated*: CCGCCATGCTTCCTTAACTA

*Mse_Aim2_F_Unannotated*: AGGCGGATGGTTTGAACTCT

*Mse_Aim2_R_Exon2*: TTGAAGCAACTTCCATCTGC

*Mse_Aim2_CDS_F*: AGTACCGGGAAATGCTGTTG

*Mse_Aim2_CDS_R*: GAGTGTGCTCCTGGCAATCT

*Mse_Gapdh_F*: CCAATGTGTCCGTCGTGGATC

*Mse_Gapdh_R*: GTTGAAGTCGCAGGAGACAAC

*Mse_Neat1_F*: TTGGGACAGTGGACGTGTGG

*Mse_Neat1_R*: TCAAGTGCCAGCAGACAGCA

### RT-PCR Validation

RT-PCR validation was completed using three biological replicates. KAPA HiFi HotStart ReadyMix PCR Kit (Kapa Biosystems) and the manufacturer’s suggested cycling protocol was used to complete the PCR reaction with the following primers:

*Mse_Denr_F1:* ATCGCGATAAAGGCTCATTG

*Mse_Denr_F2:* GCTACCTGTCCTTTTCCCCA

*Mse_Denr_R:* AACTTGGCACTGTTCTTCGT

*Mse_Arhgef7_F1:* TGTTGTTCTGGGGTTTGTGA

*Mse_Arhgef7_F2:* CTGTGTGTTGCAGGTCTACC

*Mse_Arhgef7_R:* GTGTCACCAAGGAGCTGAGG

*Mse_Ncoa7_F1:* GTGGTGGAGAAGGAAGAGCT

*Mse_Ncoa7_F2:* TTCTATTGTGCCAGGCCTGA

*Mse_Ncoa7_R:* GCATGTTTTCCAGGAGTGCA

*Mse_Ampd3_F1:* CCCTACTGTAGATGAATCCCCTTA

*Mse_Ampd3_F2:* GCTGAGCTTTGTGTCTGTGT

*Mse_Ampd3_R:* GGGGACAGTAAACAGGGACA

*Mse_Rcan1_F1:* ACTGGAGCTTCATCGACTGC

*Mse_Rcan1_F2:* GACTGAGAGAGCGAGTCGTT

*Mse_Rcan1_R:* CATCGGCTGCAGATAAGGGG

*Hu_NCOA7_F1:* TGTTCAGTGGTCTCCCGATGTCTATGG

*Hu_NCOA7_R:* GGGCCGTAGGACAGGCAGCA

*Hu_NCOA7_R2:* AGCGTGGCTACAAGTAACTGTGGTGT

*Hu_AMPD3_F1:* TATGCAAAACAGAGACCTCC

*Hu_AMPD3_R:* CACTTCAGAGATGTTCAGCT

*Hu_AMPD3_F2:* CCTGCTTGGTTTTAGAGGAT

*Hu_RCAN1_F1:* GACTGGAGCTTCATTGACTG

*Hu_RCAN1_R:* ATTCTGACTCGTTTGAAGCT

*Hu_RCAN1_F2:* TAGCGCTTTCACTGTAAGAA

Band intensities were measured for each band in each condition and sample using FIJI [Schindelin, J]. The relative abundance of each isoform was calculated using the equation to calculate PSI (PSI = inclusion/ (inclusion + exclusion)) in each condition and sample to validate the computationally derived delta PSI values. A gel extraction was completed for each band using the PCR clean-up Gel extraction kit (Machery-nagel). The PCR product was confirmed using Sanger Sequencing.

### Protein Lysate and Western Blot

Cell lysates were prepared in RIPA buffer (50 mM Tris-HCl pH 8.0, 150 mM NaCl, 1 mM EDTA, 1% (v/v) Nonidet P-40, 0.5% (w/v) sodium deoxyxholate, 0.1% (w/v) SDS) containing protease inhibitor cocktail (Roche) and quantified by the Bicinchoninic Acid Assay (BCA) assay (Thermo Fisher). Equivalent masses (15ug) of each sample were resolved by SDS-PAGE and transferred to a polyvinylidene difluoride (PVDF) membrane and Western blotted with either Aim2 (1:1,000; Cell Signaling #63660) and horseradish peroxidase-conjugated b-actin monoclonal antibody (1:5,000, Santa Cruz Biotechnology) were used as a loading control. HRP-conjugated goat anti-rabbit (1:1,500, Biorad) secondary antibodies were used. Image J (*90*) was used for quantification of Western blots.

### Alternative splicing quantification

After the identification of each alternative splicing event, JuncBASE counts reads supporting the inclusion and exclusion isoform of each event. Isoform abundances are then calculated by dividing the read counts for the isoform by the length of the isoform. Ψ-values for each splicing event are derived from the isoform abundances:

PSI Formula:

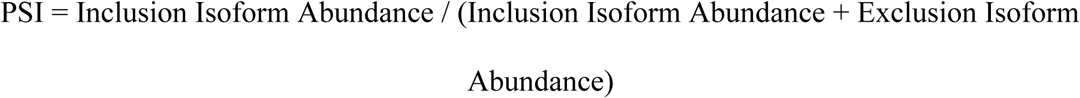

### Statistical Analysis

Error bars represent the standard deviation of biological triplicates. Student’s t-tests were performed using GraphPad Prism. Asterisks indicate statistically significant differences between mouse lines (*p> 0.05, **p> 0.01, ***p>0.005).

### Illumina RNA Sequencing (Human)

RNA-seq libraries were prepared with the Illumina TruSeq RNA Sample Preparation kit (Illumina) according to the manufacturer’s protocol. Libraries were validated on an Agilent Bioanalyzer 2100. Indexed libraries were equimolarly pooled and sequenced on a SE50 (single-end 50base pair) Illumina HiSeq2500 lane, which yielded an average of about 30 × 10^6^reads/sample.

### Illumina RNA Sequencing (Mouse)

For generation of RNASequencing libraries, RNA was isolated as described above and the RNA integrity was tested with a BioAnalyzer (Agilent Technologies) or FragmentAnalyzer (Advanced Analytical). For RNASequencing target RIN score of input RNA (500-1000ng) usually had a minimum RIN score of 8. RNASequencing libraries were prepared with TruSeq stranded RNA sample preparation kits (Illumina), depletion of ribosomal RNA was performed by positive selection of polyA+ RNA. Sequencing was performed on Illumina HighSeq or NextSeq machines. RNA-seq 50bp reads were aligned to the mouse genome (assembly GRCm38/mm10) using TopHat (*91*). The Gencode M13 gtf was used as the input annotation. Differential gene expression specific analyses were conducted with the DESeq (*92*) R package. Specifically, DESeq was used to normalize gene counts, calculate fold change in gene expression, estimate p-values and adjusted p-values for change in gene expression values, and to perform a variance stabilizing transformation on read counts to make them amenable to plotting. Data was submitted to GEO GSE141754.

### Nanopore Direct RNA Sequencing

#### Total RNA extraction

Total RNA was extracted according to Workman *et al*. (*93*). 5 x 10^7^ frozen macrophages were resuspended in three ml of TRI-Reagent (Invitrogen AM9738) and vortexed for 5 min. The mixture was incubated at RT for 5 minutes, transferred to 1.5 mL tubes and spun down to remove debris. Supernatant was transferred to fresh tubes and chloroform extracted. The aqueous portion was mixed with an equal volume of isopropanol, incubated for 15 min at RT and centrifuged at 12,000 g at 4C. Pellet was washed twice with 75% ethanol, air dried and resuspended in nuclease free water.

#### Poly(A) RNA isolation

One hundred μg aliquots of total RNA preparation were brought to 100 μl in nuclease free water and poly-A selected using NEXTflex Poly(A) Beads (BIOO Scientific Cat#NOVA-512980) according to the manufacturer’s instructions. The resulting poly-A RNA solution was stored at −80°C.

#### Library preparation and MinION run

A native RNA sequencing sequencing library was prepared following the ONT SQK-RNA001 using Superscript IV (Thermo Fisher) for the reverse transcriptase step. Sequencing was performed using ONT R9.4 flow cells and the standard MinKNOW protocol.

#### Basecalling and sequence alignments

ONT albacore version 2.1.0 was used to baseball Nanopore direct RNA raw signal. We used minimap2 (*94*) with default parameters to align reads to the mm10 mouse genome reference. Following alignment, we used SAMtools (*95*) to filter-out reads with mapping quality (MAPQ) less than 30.

### Alignment of Paired-end Mouse RNA-seq data

Bowtie2-build v2.3.1 (*96*) was used to build the index files from GRCm38.p6 mouse (mm10 assembly) genome sequence obtained from Gencode. The index files were then used for completing paired-end alignment of each sample using TopHat2 v2.1.1 (*97*) with parameters: *--segment-length 20, --library-typ fr-firststrand, --no-coverage-search*.

### Identification of Splicing Events and Calculating PSI

Human monocyte-derived macrophage +/− LPS and Mouse bone marrow-derived macrophage +/− LPS were each run through JuncBASE v1.2 (*98*) to calculate percent spliced in (PSI) values and identify splicing events. The JuncBASE parameters used for the identification of splicing events and calculation of PSI in Human monocyte derived macrophage +/− LPS are: *-c 1.0 -j [introns from Gencode v24 (hg19 assembly)* (*99*) *--jcn_seq_len 88*. The JuncBASE parameters used for the identification of splicing events and calculation of PSI in Mouse bone marrow-derived macrophage +/− LPS are: *-c 1*.*0 -j [introns from Gencode M18 (mm10 assembly)*(*99*) *--jcn_seq_len 88*.

### Identification of High-Confidence Isoforms from Nanopore Data

FLAIR (full length alternative isoform analysis of RNA) (*100*) was used to assemble the high-confidence isoforms from native RNA sequencing of Mouse BMDM +/− 6hr LPS. FLAIR modules align, correct, and collapse were used for the assembly. Corresponding short read data was used when running the correct module in order to help increase splice-site accuracy. Putative promoter regions were obtained using ATAC-seq data from Smale *et al*. and Atianand *et al*. (*101, 102*) converted to mm10 coordinates using liftOver (*103*), and used when running the collapse module.

### Differential Splicing Analysis

Differential splicing analysis was completed using DRIMSeq v1.10.1 (*104*) and the compareSampleSets.py script within JuncBASE. CompareSampleSets.py applies the statistical t-test and DRIMSeq applies the framework of the Dirichlet-multinomial distribution for differential analysis. Each tool was used to apply the respective statistical method in order to determine significant differentially spliced events between control (-LPS) and LPS (+LPS) conditions. The AS_exclusion_inclusion_counts_lenNorm.txt JuncBASE output table from the identification and quantification analysis of each experiment was used as the input table for both compareSampleSets.py and DRIMSeq.

For all experiments, compareSampleSets.py was run using parameters: *--mt_correction BH --which_test t-test --thresh 10 --delta_thresh 5*.*0*. The following parameters were used for the differential splicing analysis of data from Human monocyte-derived macrophage +/− LPS with DRIMSeq: *min_samps_gene_expr = 8, min_samps_feature_expr = 4, min_gene_expr = 10, min_feature_expr = 0*. The following parameters were used for the differential splicing analysis of data from Mouse bone marrow-derived macrophage +/− LPS with DRIMSeq: *min_samps_gene_expr = 6, min_samps_feature_expr = 3, min_gene_expr = 10, min_feature_expr = 0*. Following differential splicing analysis using each tool, genes with significant differential splicing events were filtered for using thresholds of a corrected/adjusted p-value ≤ 0.25 and a |Δ PSI| ≥ 10. Within each category of event type, the union of genes with significant events identified using compareSampleSets.py and DRIMSeq within each experiment was used for further comparison. Novel intron retention events, annotated with a “N,” were removed for further analyses.

### Creating merged reference annotation files and incorporating nanopore

The *isoforms*.*gtf* output file from FLAIR collapse was combined with the Gencode M18 (mm10 assembly) basic annotation using cuffmerge from Cufflinks v2.2.1 (*105*) with parameter: -s GRCm38.p6.genome.fa. Similarly, the *isoforms*.*gtf* output file was combined with the Gencode M18 (mm10 assembly) comprehensive annotation with parameter: -s GRCm38.p6.genome.fa. The resulting comprehensive annotation output file was used to generate an intron coordinate file for the identification of splicing events and calculating PSI of splicing events found in Mouse Bone-marrow derived macrophage +/− 6hr LPS using JuncBASE with parameters: *-c 1*.*0, -j [intron coordinates from merged comprehensive annotation], --jcn_seq_len 238*. Parameters used for finding significantly differentially spliced events using compareSampleSets.py from JuncBASE are: *--mt_correction BH --which_test t-test --thresh 10 --delta_thresh 5*.*0*. Parameters used for finding significantly differentially spliced events using DRIMSeq are: *min_samps_gene_expr = 6, min_samps_feature_expr = 3, min_gene_expr = 10, min_feature_expr = 0*.

### Gene Expression Analysis

DESeq2 v1.22.2 (*106*) was used to create counts tables and complete differential gene expression analysis on RNA-seq data from Human monocyte-derived macrophage +/− 18hr LPS and Mouse BMDM +/− 6hr LPS experiments. The sample conditions used were “control” and “LPS.” Data was plotted using ggplot2 v3.1.1 (*107*). Significance thresholds were set to |log2FC| ≥ 2 and adjusted p-value ≤ 0.05. The list of genes with significant AFE events were then highlighted on the appropriate graphs.

### Creating and comparing gene lists

For each experiment, a table with the union of significant events found using DRIMSeq and compareSampleSets.py was created. A list of genes with significant events was generated for each experiment using this table. BioVenn (*108*) and DrawVenn (*94*) were then used to remove duplicate gene names and compare the lists of genes to find unique and common genes between experiments.

### Differential chromatin accessibility

Raw ATAC-seq fastq sequence files were published in Tong *et al*. (*102*) and pulled from the GEO accession number GSE67357. A bowtie2 (*96*) index file was created from the GENCODE mm10 version M18 genome annotation file and the untreated and LPS treated ATAC-seq reads were aligned using the created index file with the default bowtie2 parameters. Peaks were then called separately by treatment type on untreated and treated samples using the ENCODE published ATAC-seq peak calling pipeline (https://github.com/ENCODE-DCC/atac-seq-pipeline) using the aligned reads as sequence input. Parameters that were used followed the basic JSON input file template, using an IDR threshold of 0.05. Peaks from both conditions were then merged using bedtools merge (*109*) if the tail ends were less than 10bp away from each other, in order to create a set of consensus peaks from both conditions. A GFF file was created from the merged peaks, assigning a unique ID to each peak. This GFF file was provided to HTSeq-count (*110*) along with the aligned reads for each condition in each replicate to count reads aligning to the unique peaks. The read count matrix was provided to DESeq2 (*106*) to call differential peaks. All peaks were considered significant if log2FC ≥ 0.8 and p-value ≤ 0.15.

### Differential transcription factor binding

ChIP-seq fastq sequencing files for the NF-κB subunit p65 and interferon transcription factor Irf3 were downloaded from the GEO accession number GSE67357 published by Tong *et al*. (*102*). ChIP-seq samples for input control, untreated, and LPS treatment were aligned using bowtie2 (*96*) to the mm10 version M18 mouse genome annotation with default parameters. Peaks were separately called between untreated and treated conditions using the ENCODE published ChIP-seq peak calling pipeline (https://github.com/ENCODE-DCC/chip-seq-pipeline2) from the aligned reads. The aligned input control reads were input as genomic background to account for noise in ChIP-seq experiments. The basic JSON input template file was used, using an IDR threshold of 0.05. Differential peak analysis was done using the HOMER suite designed for ChIP-seq data (*111*). Consensus peaks from both conditions were merged using mergePeaks within HOMER, reporting the direct overlap between peaks. Tag directories were created to count reads for each aligned sequence file with TagDirectory. The merged consensus peaks were then annotated for raw read counts using the tag directories for each replicate and condition with the annotatePeaks.pl tool. Annotated consensus peaks were provided to getDiffExpression.pl, normalizing to total read counts. Peaks were considered significant if they had a p-value ≤0.25and log2FC ≥ 1.

### AFE event overlap

To identify differential transcription factor binding and chromatin remodeling at the promoters of the observed alternative first exon events, the coordinates of the alternate first exon were determined from the statistical testing results. Significant (p-value ≤ 0.05) alternative first exon events were first filtered out from all results. For all significant results, if the inclusion exon had a Δ PSI *≥* 10, the inclusion exon coordinates from the JuncBASE table were used as the coordinates for that splicing event. If the inclusion exon had a Δ PSI ≥ −10, all other inclusion exons for that splicing event from the statistical testing (DRIMSeq or t-test) were considered, and any inclusion exon with Δ PSI > 10 was used. Redundant events with the same exon coordinates were then filtered out, leading to a final set of 77 exon coordinates. The coordinates were then extended to include 10kbp upstream of the exon. Overlap of differential chromatin accessibility and different transcription factor binding was done using bedtools intersect (*109*) with the significant differential peak coordinates and the alternative first exon 10kbp upstream region, returning the coordinates of the exon that show differential chromatin accessibility or transcription factor binding.

---

**Supplemental Fig 1:**
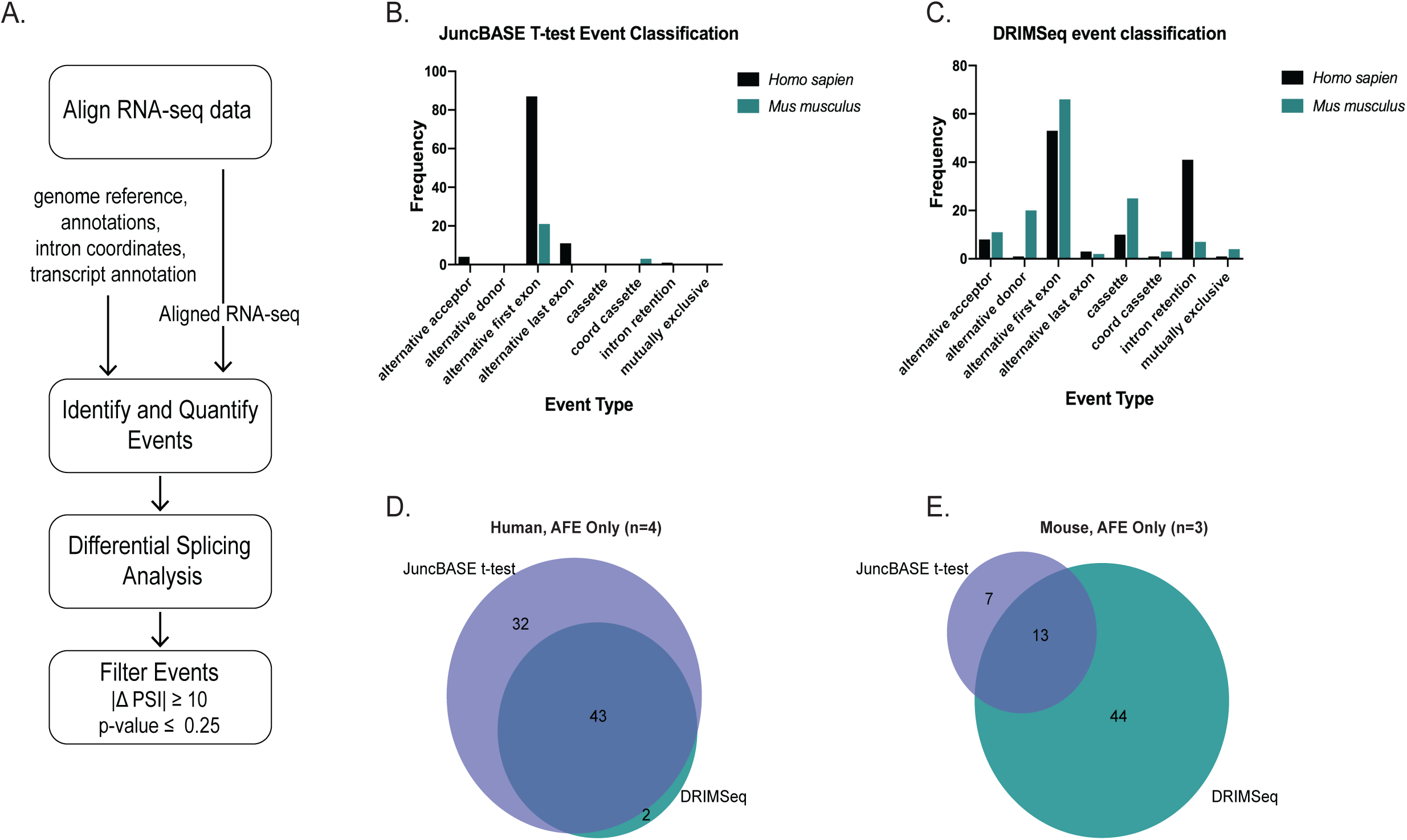
Computational Pipeline and comparison of t-test and DRIMSeq alternative splicing events. (A) Bioinformatic pipeline for Human and Mouse RNA-seq data. (B) Alternative splicing event type classification of significant differential splicing events (|ΔPSI| ≥ 10 and corrected p-value ≤ 0.25) in Human and Mouse macrophages +/− LPS as identified and quantified using the t-test with JuncBASE. The AFE event is the most predominant in both human and mouse samples. (C) AS splicing event type classification of significant differential splicing events (|ΔPSI| ≥ 10 and corrected p-value ≤ 0.25) in human and mouse macrophages +/− LPS as quantified using the Dirichlet-multinomial framework applied by DRIMSeq. The AFE event is the most predominant in both human and mouse samples. (D)A comparison of genes found to have significant AFE events following +/− LPS stimulation of Human monocyte-derived macrophage cells by the JuncBASE t-test and DRIMSeq. There were 32 genes unique to the analysis that applied the JuncBASE t-test, 2 genes unique to the analysis that applied DRIMSeq, and 43 genes in common between the two sets of analyses. (E) A comparison of genes found to have significant AFE events following +/− LPS stimulation of Mouse bone marrow-derived macrophage cells by the JuncBASE t-test and DRIMSeq. There were 7 genes unique to the analysis that applied the JuncBASE t-test, 44 genes unique to the analysis that applied DRIMSeq, and 13 genes in common between the two sets of analyses.

**Supplemental Fig. 2:**
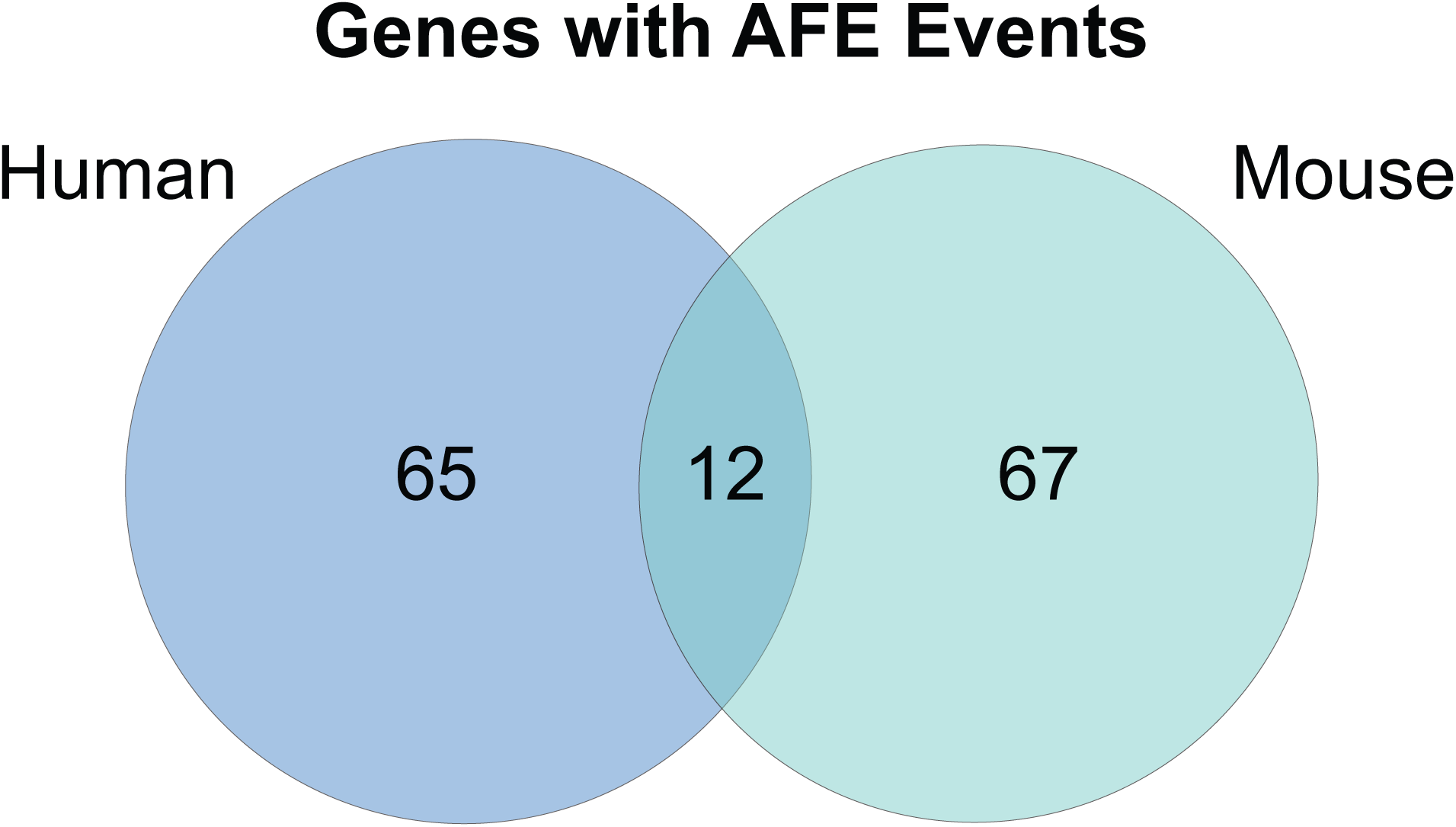
AFE conservation in mouse and human. Venn diagram denoting unique and overlapping genes with significant alternative first exon (AFE) switches in human and mouse.

**Supplemental Fig. 3:**
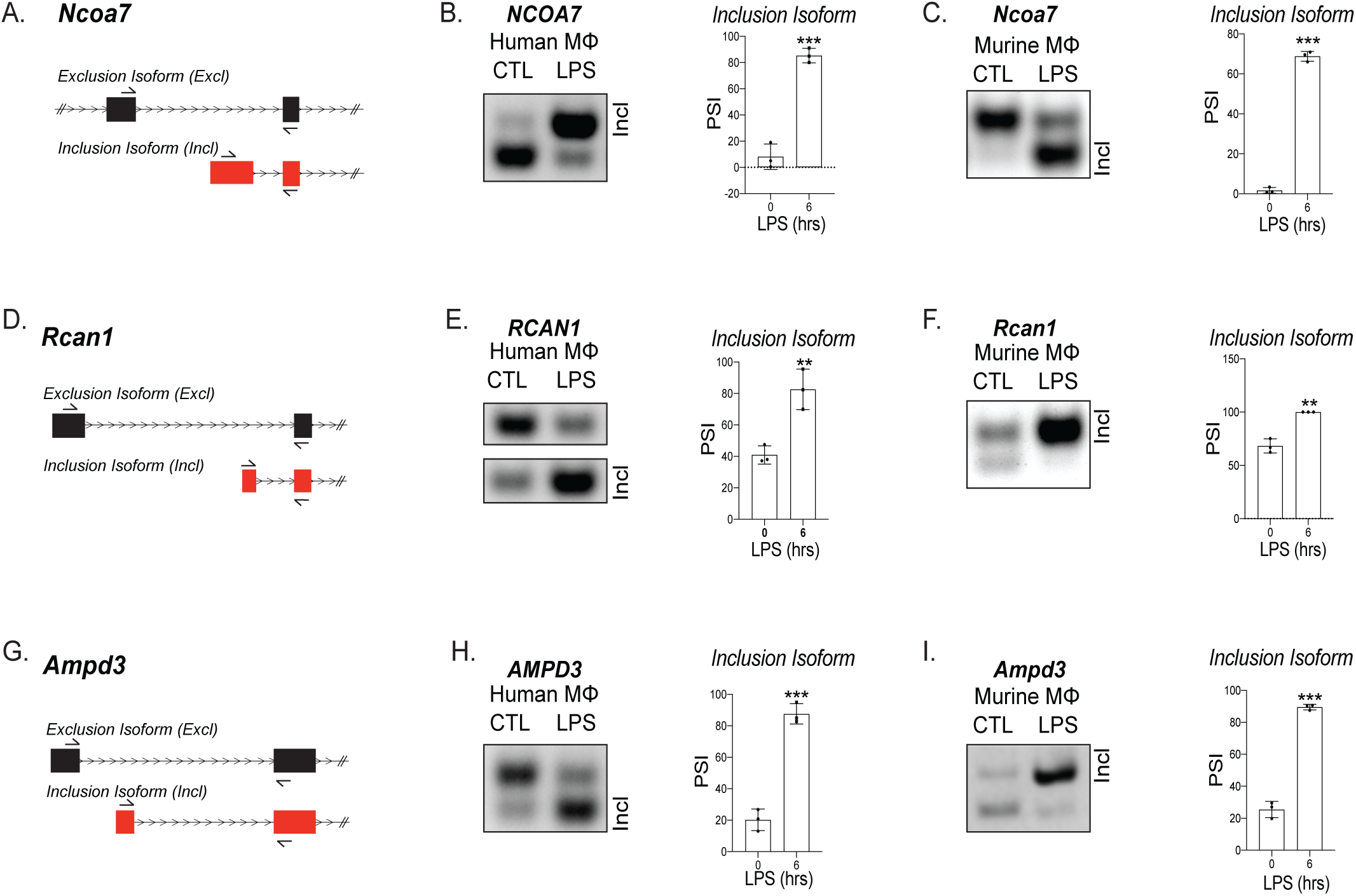
Validation of mouse and human alternative first exon events. (A) mRNA transcript diagram of the exclusion and inclusion isoform of Ncoa7 for human and murine macrophages. RT-PCR of human (B) and murine (C) macorphages, Rcan1 (D-F) and Ampd3 (G-I) with PSI calculation.

**Supplemental Fig. 4:**
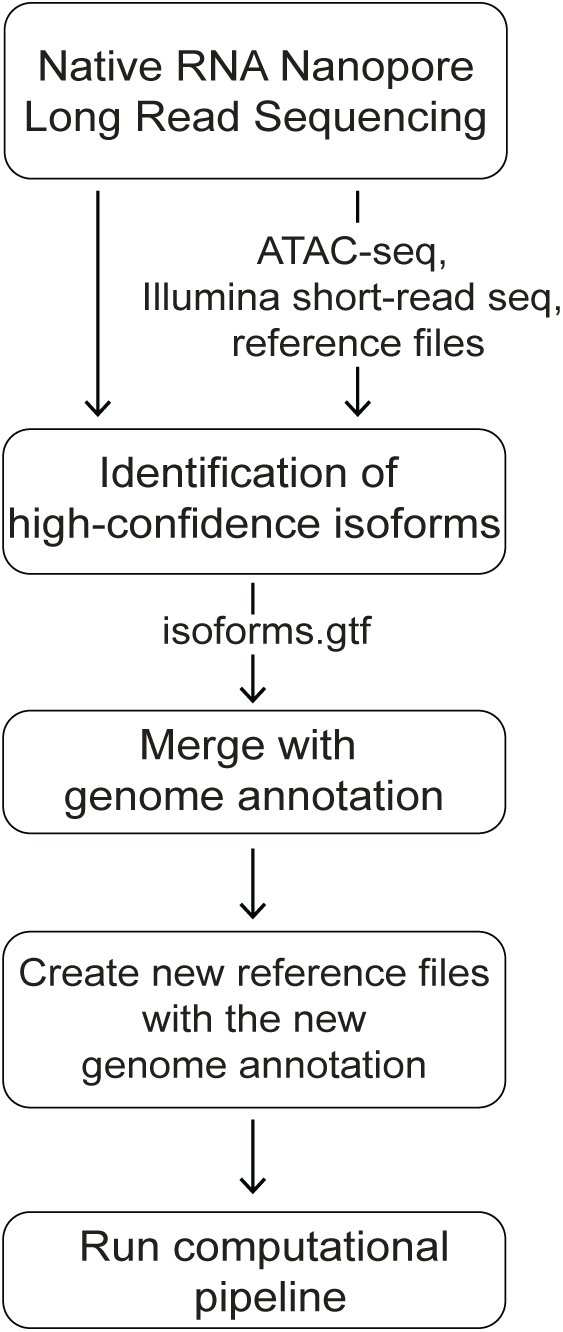
Bioinformatic pipeline. Schematic describing the steps taken to analyze alternative splicing of native RNA nanopore sequencing.

**Supplemental Fig. 5:**
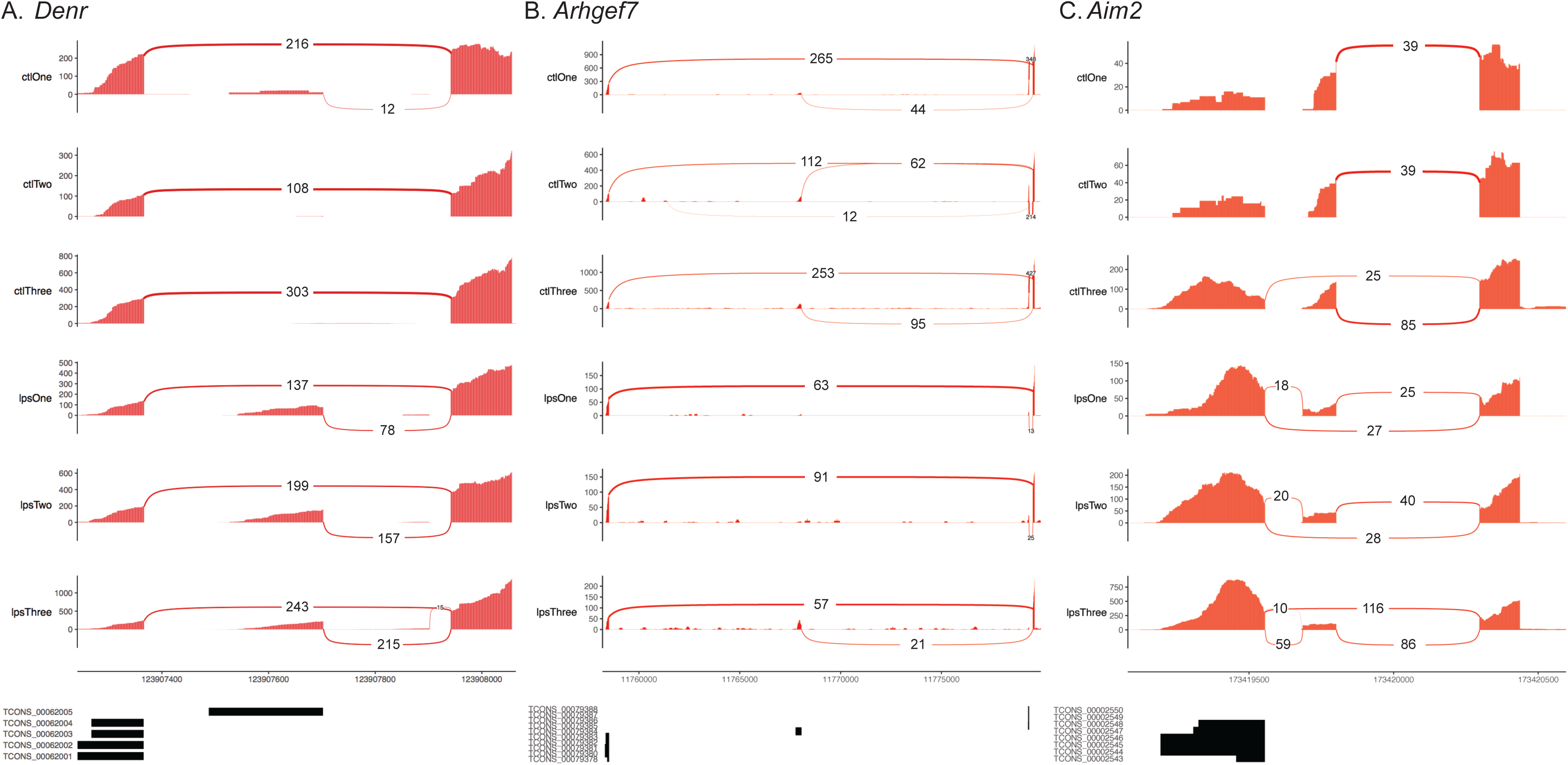
Validated unannotated alternative first exon isoforms. Sashimi plots showing AFE usage in (A) Denr, (B) Arhgef7, and (C) Aim2 involving novel isoforms identified using nanopore sequencing.

**Supplemental Fig 6:**
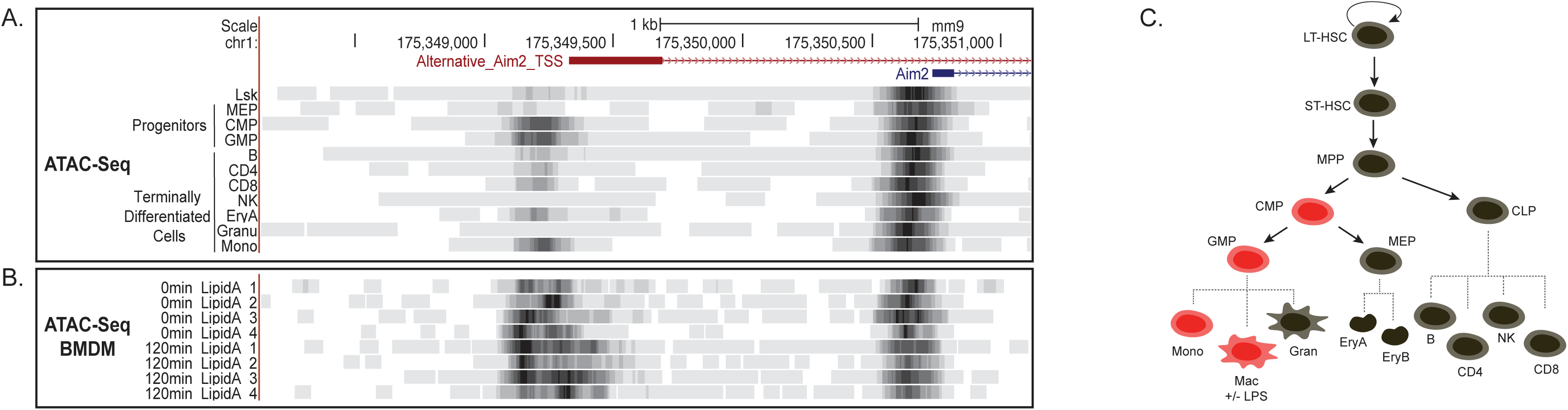
Chromatin remodeling not a mechanism driving novel Aim2 isoform. (A) mm9 genome browser shot between chr1:175,348,283-175,351,422 indicating ATAC-seq of cells from hematopoietic tree. (B) mm9 genome browser shot between chr1:175,348,283-175,351,422 indicating ATAC-seq of BMDMs +/− Lipid A for 2 hrs. (C) Schematic summarizing promoter accessibility of novel Aim2 isoform, red=accessible and black=not accessible.

**Supplemental Fig 7:**
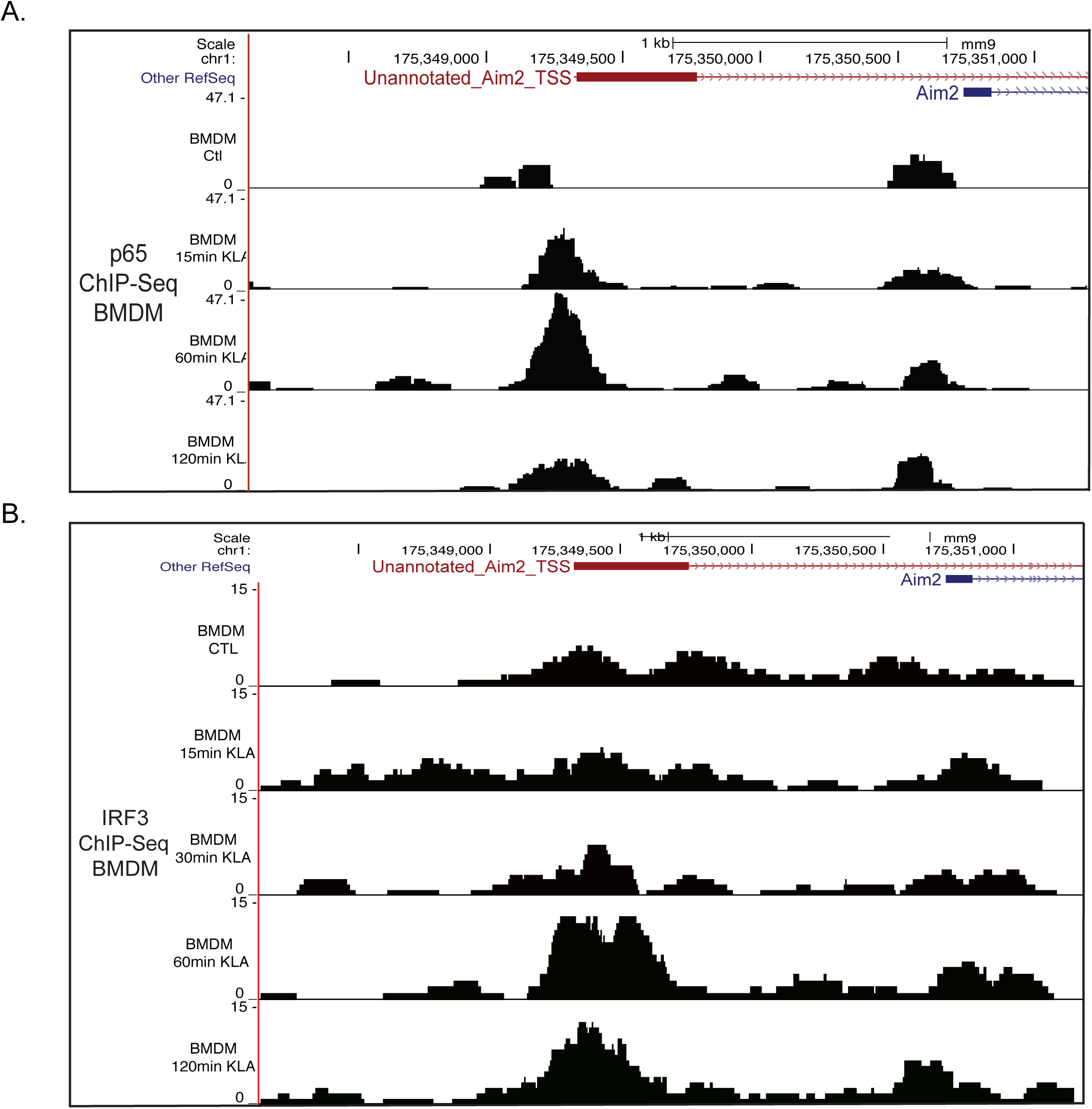
Novel Aim2 isoform driven by IRF3 and p65 transcription factors. mm9 genome browser shot between chr1:175,348,283-175,351,422 for ChIP-seq of p65 binding (A) and IRF3 binding (B).

**Supplemental Fig 8:**
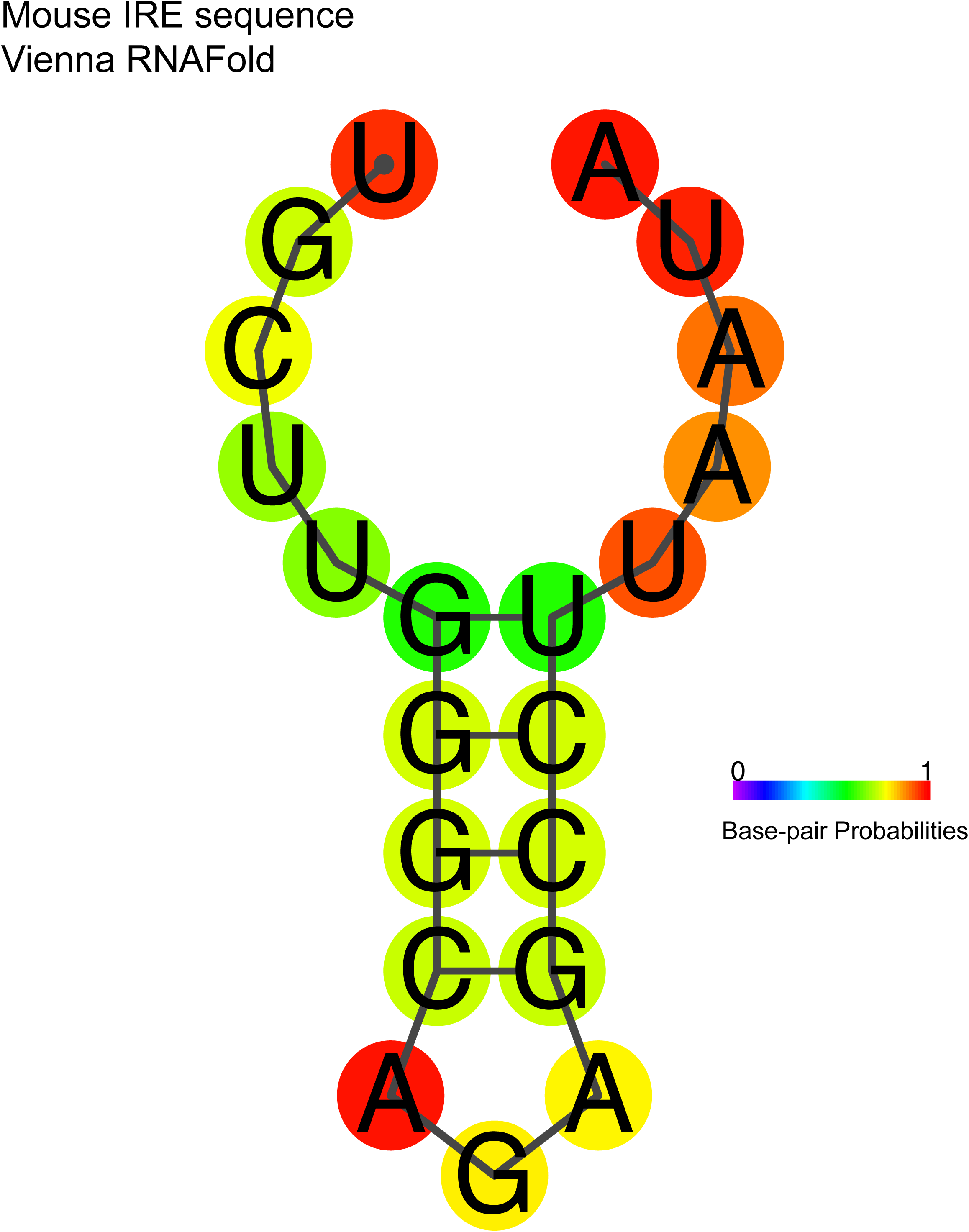
IRE hairpin structure predicted. Using the Vienna RNAFold package, the IRE motif is folded into a hairpin, with strong base-pairing.

